## Supplementary File for "Transgenerational effects of early life stress on the fecal microbiota in mice"

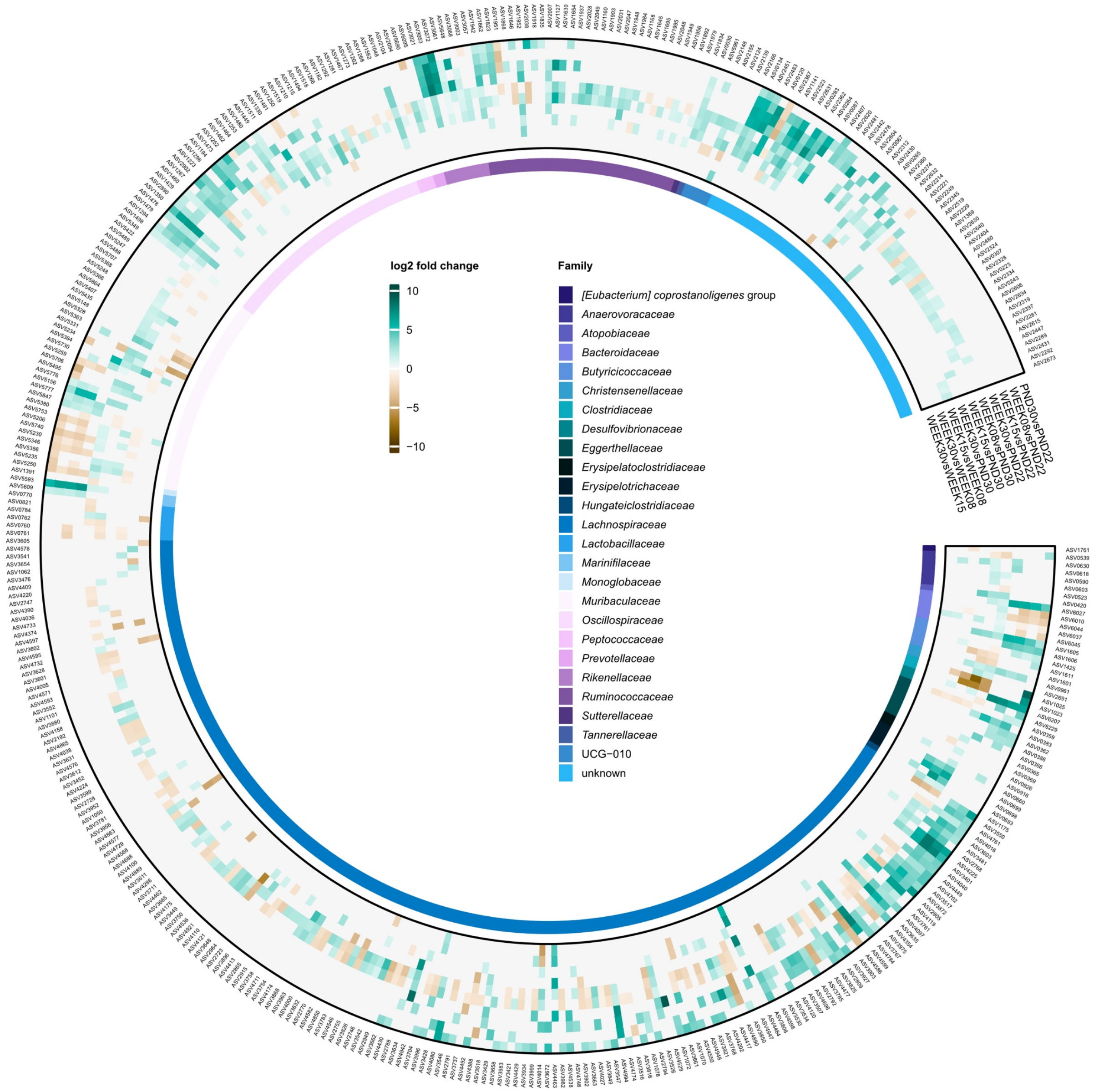


**Supplementary Figure 1: Effect of age on relative abundance of ASVs of F1 control mice fecal microbiota.** $Log_2$ fold-change in specific ASVs significantly ($p < 0.05$, FDR-adjusted) decreased or increased across life span. Visualization of pairwise comparison of different ages. Taxonomic information is indicated at family level. Significances were calculated using $log_2$ transformed abundance counts and generalized mixed effect models with FDR correction. PND: postnatal day

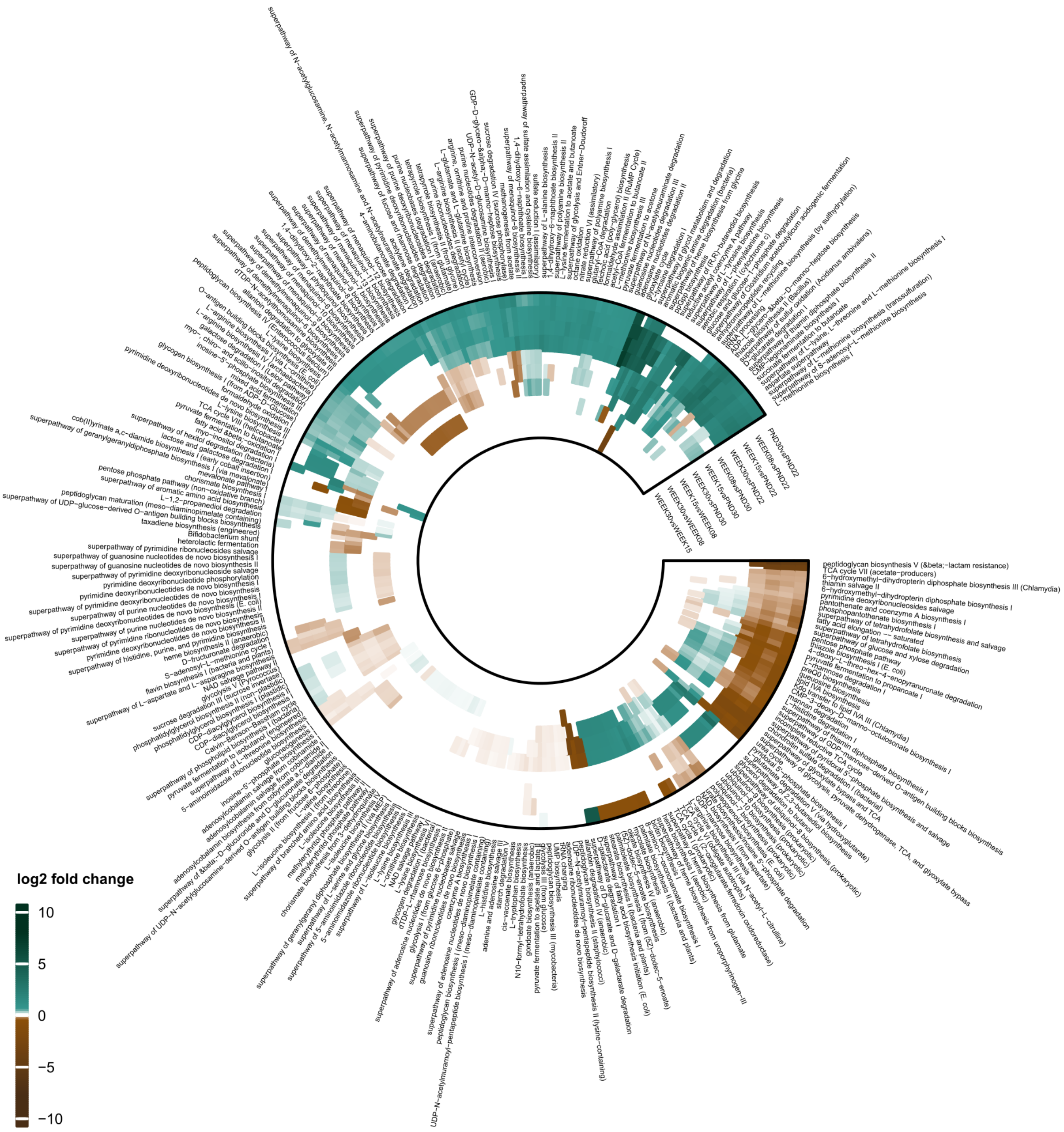


**Supplementary Figure 2: Effect of age on relative abundance of predicted fecal bacterial metabolic pathways of control mice fecal microbiota.** $Log_2$ fold-change in predicted fecal bacterial metabolic pathways significantly ($p < 0.05$, FDR-adjusted) increased or decreased across life span. Visualization of pairwise comparison of different ages. Significances were calculated using $log_2$ transformed abundance counts and generalized mixed effect models with FDR correction. PND: postnatal day

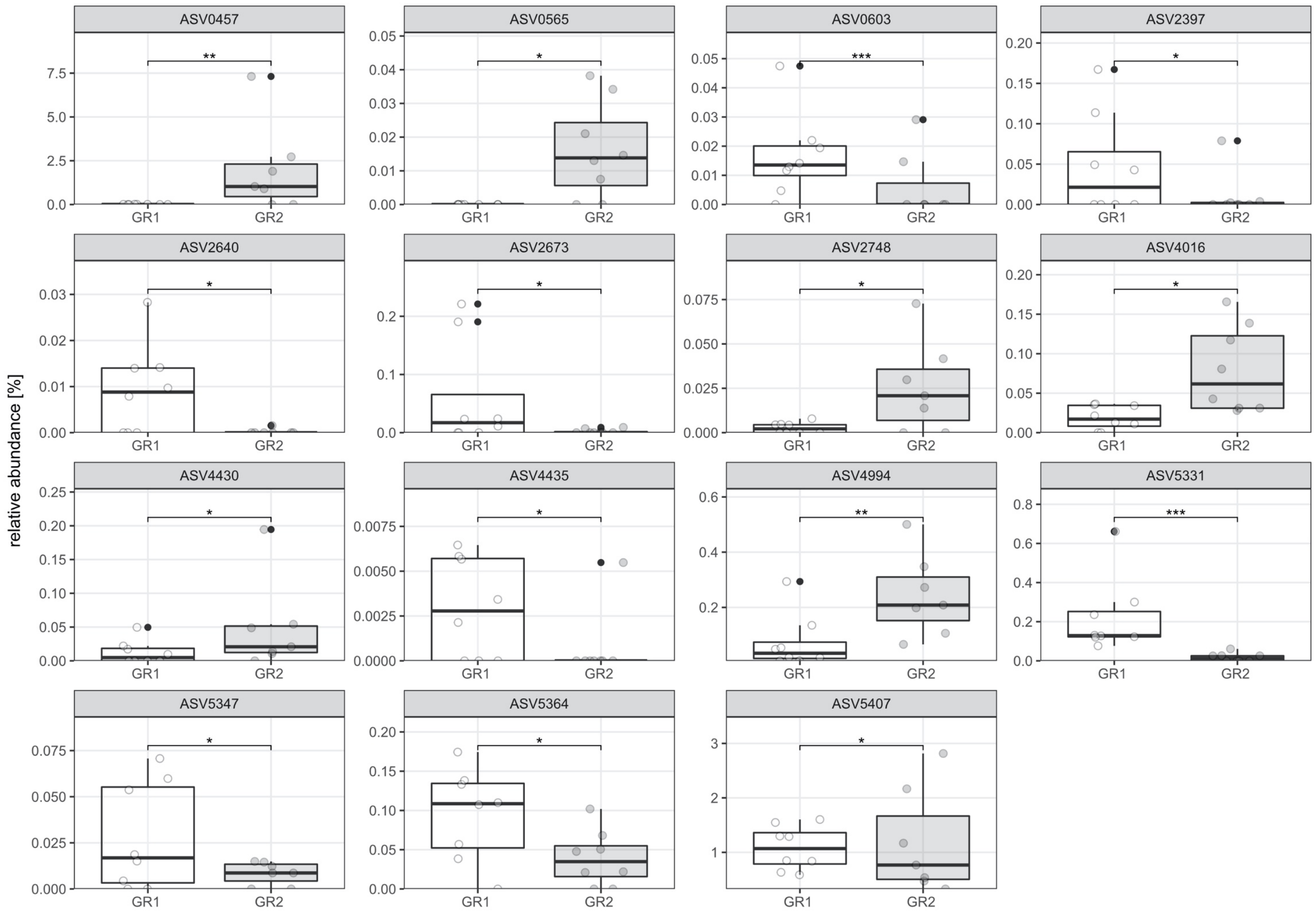


**Supplementary Figure 3: Effect of temporal succession of behavioral phenotyping and breeding on relative abundance of ASVs in 30-week-old control mice fecal microbiota.** Comparison of phenotyping group 1 (GR1; behavioral phenotyping before breeding) *versus* group 2 (GR2; breeding before behavioral phenotyping) microbiota**.** Significantly differentially abundant ASVs are displayed. Boxplot with box elements showing upper and lower quantile and median. Whiskers extend from the hinge to ± 1.5 times the interquartile range or the highest/lowest value. Outliers are indicated as black points, while white and grey points indicate individual microbiota. Significances were calculated using generalized mixed effect models with FDR correction. * $p < 0.05$; ** $p < 0.01$; *** $p < 0.001$

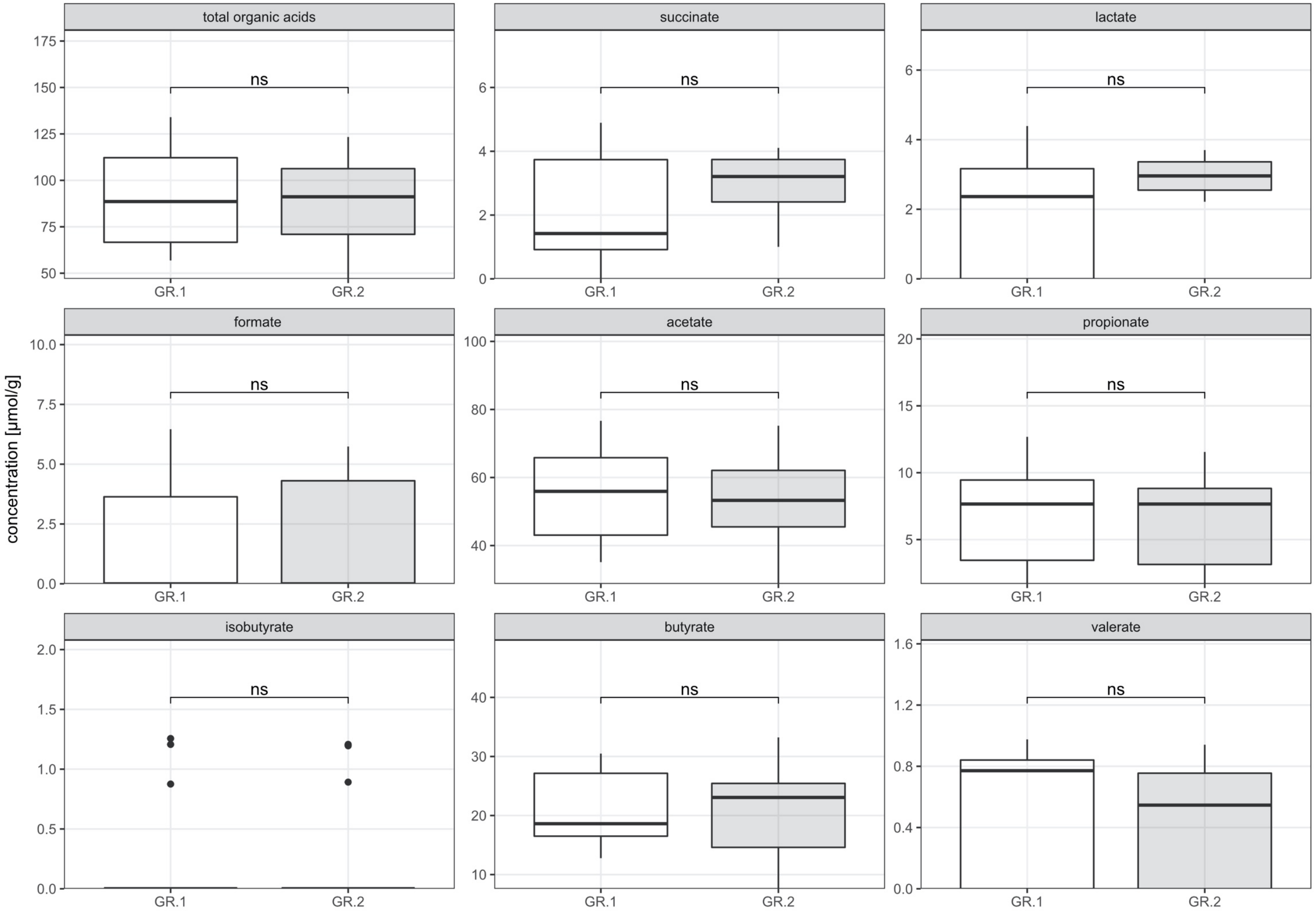


**Supplementary Figure 4: Effect of temporal succession of behavioral phenotyping and breeding on cecal total and specific organic acid concentrations of 30-week-old control mice.** Comparison of phenotyping group 1 (GR.1; behavioral phenotyping before breeding) *versus* group 2 (GR.2; breeding before behavioral phenotyping) microbiota is displayed**.** Metabolite concentration is depicted per gram cecal wet weight. Boxplot with box elements showing upper and lower quantile and median. Whiskers extend from the hinge to ± 1.5 times the interquartile range or the highest/lowest value. Outliers are indicated as black points. Significances were calculated using generalized mixed effect models with FDR correction. ns: not significant

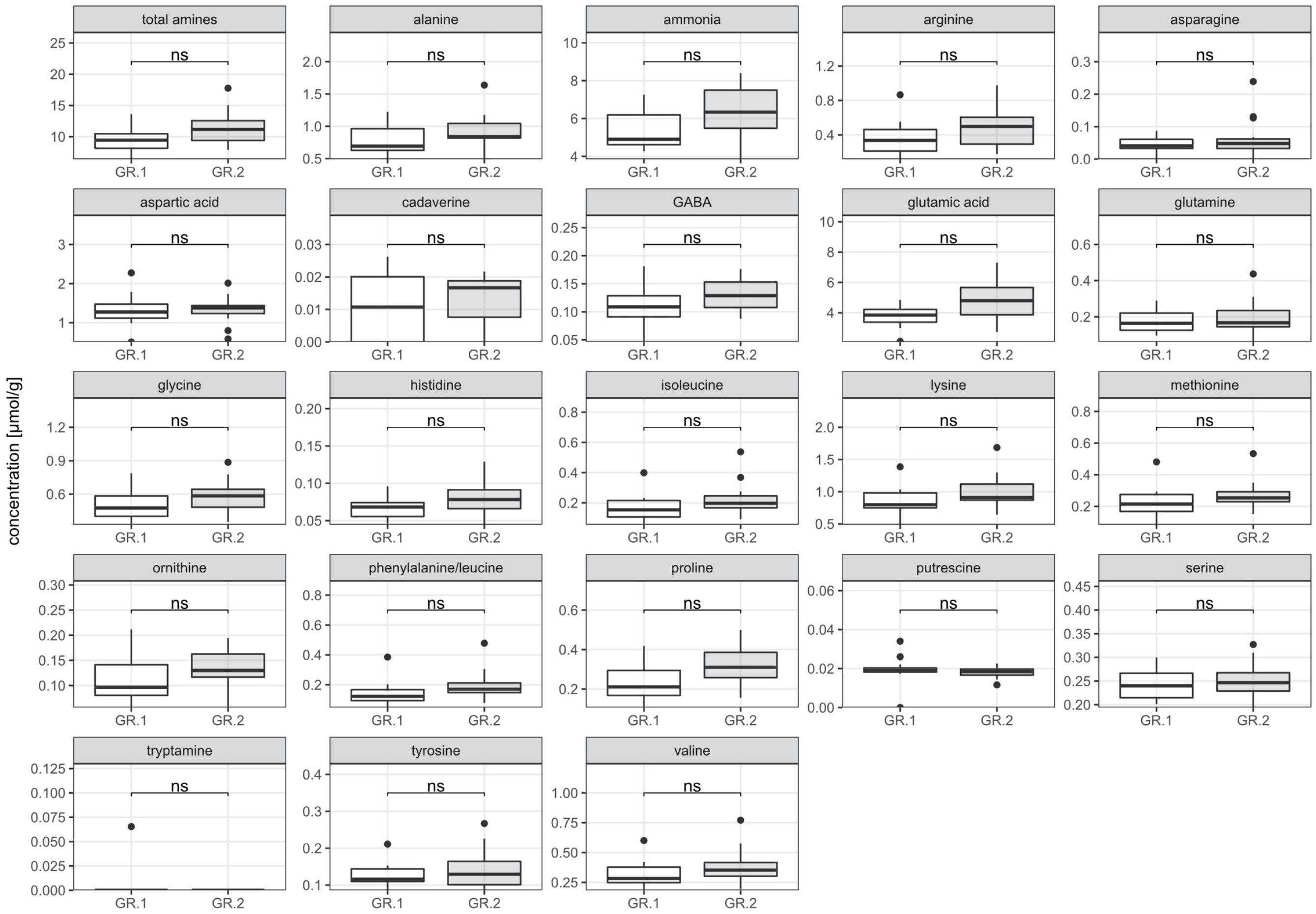


**Supplementary Figure 5: Effect of temporal succession of behavioral phenotyping and breeding on cecal amine, amino acid, and ammonia concentrations of 30-week-old control mice.** Comparison of phenotyping group 1 (GR.1; behavioral phenotyping before breeding) *versus* group 2 (GR.2; breeding before behavioral phenotyping) microbiota is displayed**.** Metabolite concentration is depicted per gram cecal wet weight. Boxplot with box elements showing upper and lower quantile and median. Whiskers extend from the hinge to ± 1.5 times the interquartile range or the highest/lowest value. Outliers are indicated as black points. Significances were calculated using generalized mixed effect models with FDR correction. ns: not significant

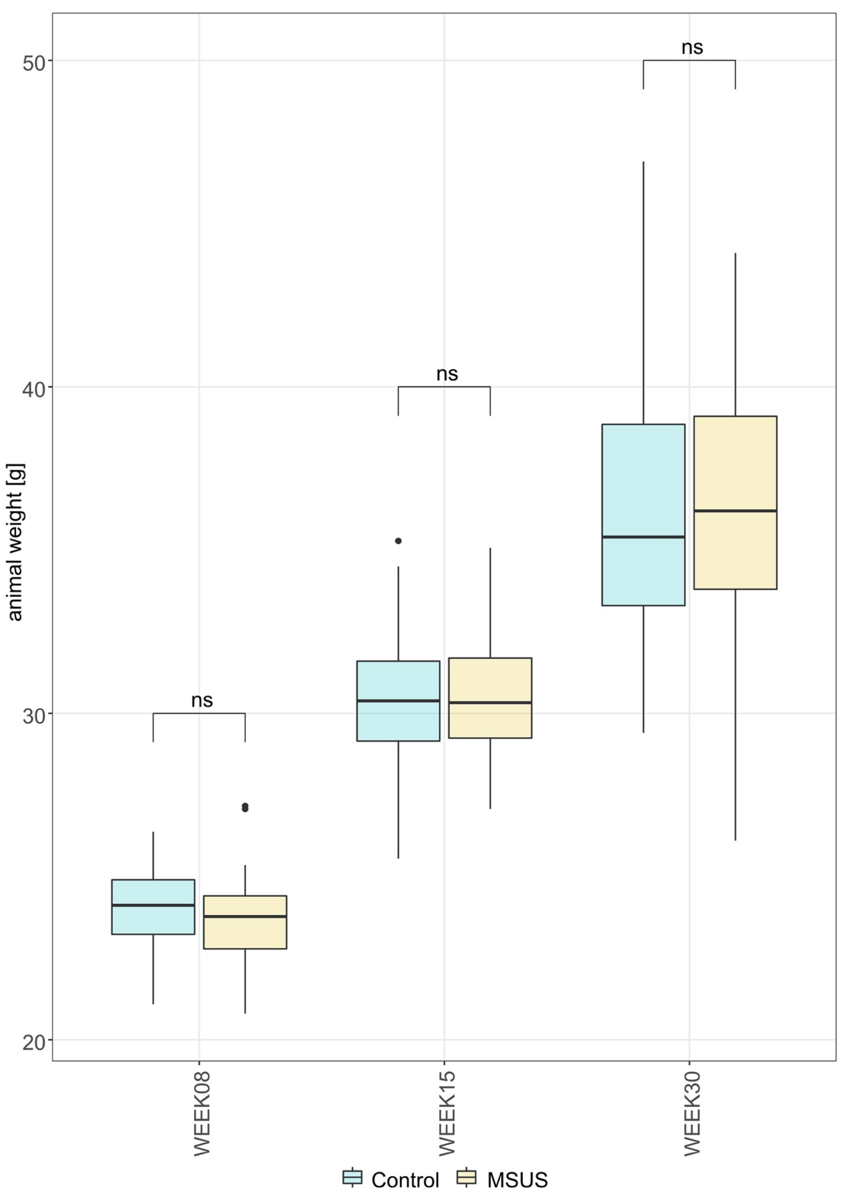


**Supplementary Figure 6: Effect of MSUS paradigm on animal weight of F1 mice across life span.** Comparison of MSUS *versus* control mice. Boxplot with box elements showing upper and lower quantile and median. Whiskers extend from the hinge to ± 1.5 times the interquartile range or the highest/lowest value. Outliers are indicated as black points. Significances were calculated using generalized mixed effect models with FDR correction. ns: not significant

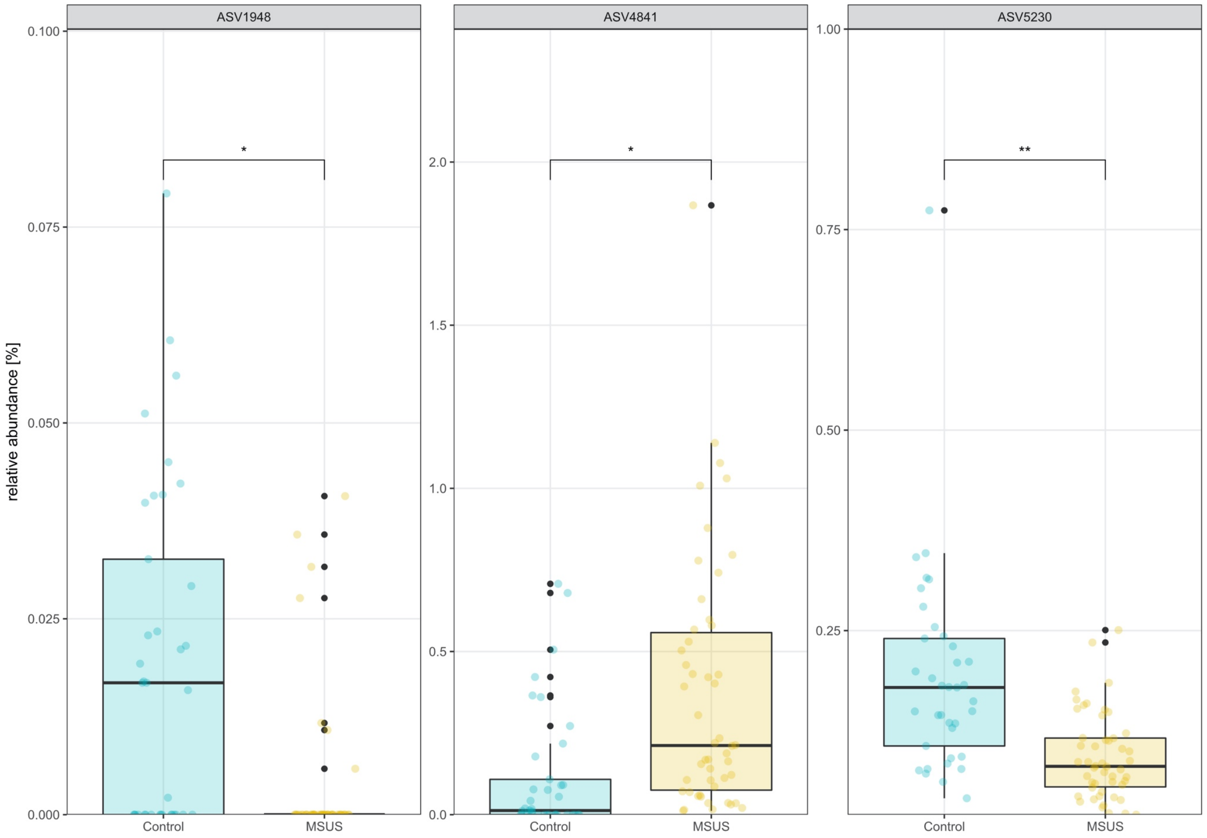


**Supplementary Figure 7: Effect of MSUS paradigm on relative abundance of ASVs in 8-week-old F1 mice fecal microbiota.** Significantly differentially abundant ASVs between MSUS *versus* control mice are displayed. Boxplot with box elements showing upper and lower quantile and median. Whiskers extend from the hinge to ± 1.5 times the interquartile range or the highest/lowest value. Outliers are indicated as black points, while colored points indicate individual microbiota. Significances were calculated using generalized mixed effect models with FDR correction. * $p < 0.05$; ** $p < 0.01$

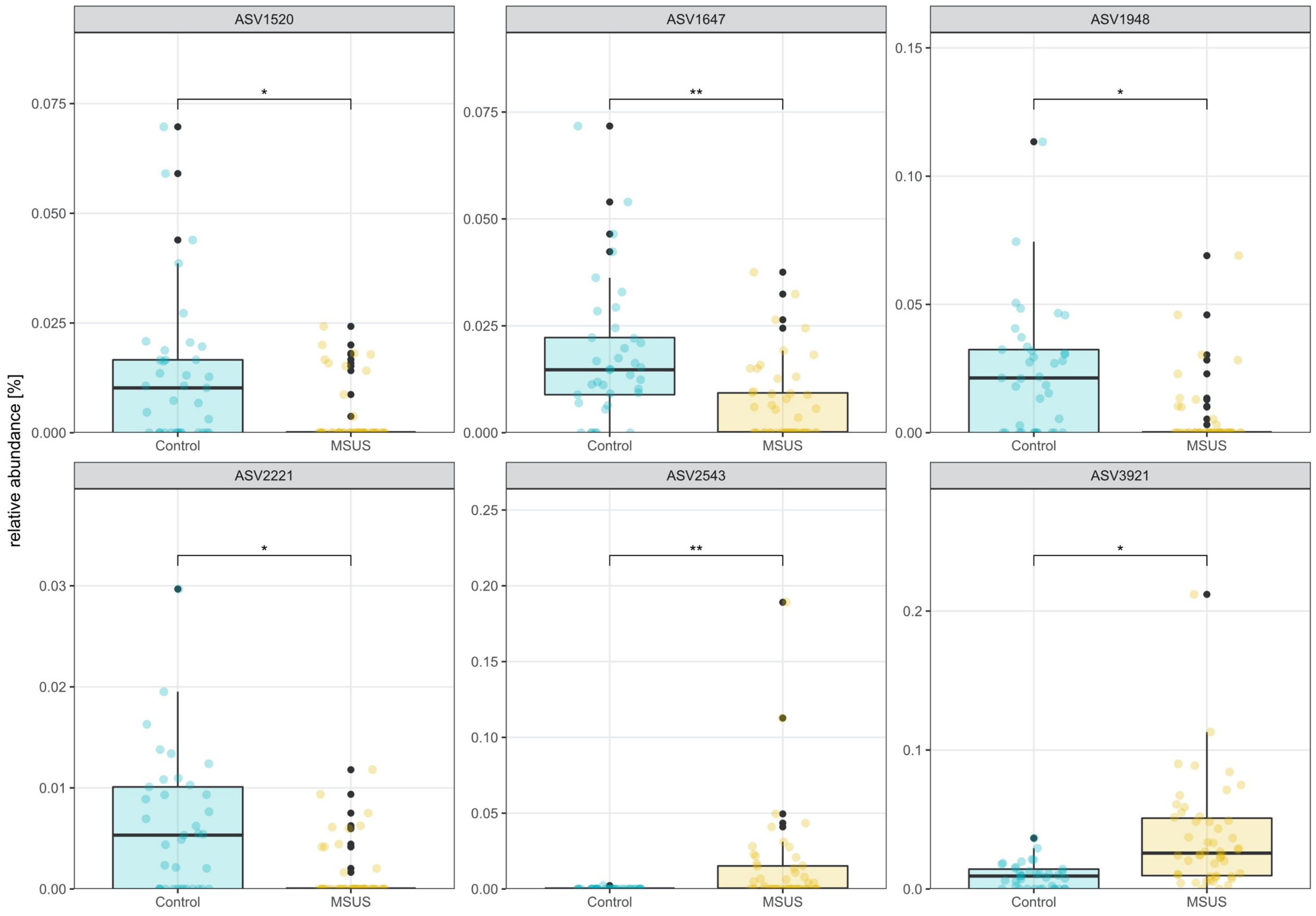


**Supplementary Figure 8: Effect of MSUS paradigm on relative abundance of ASVs in 15-week-old F1 mice fecal microbiota.** Significantly differentially abundant ASVs between MSUS *versus* control mice are displayed. Boxplot with box elements showing upper and lower quantile and median. Whiskers extend from the hinge to ± 1.5 times the interquartile range or the highest/lowest value. Outliers are indicated as black points, while colored points indicate individual microbiota. Significances were calculated using generalized mixed effect models with FDR correction. * $p < 0.05$; ** $p < 0.01$

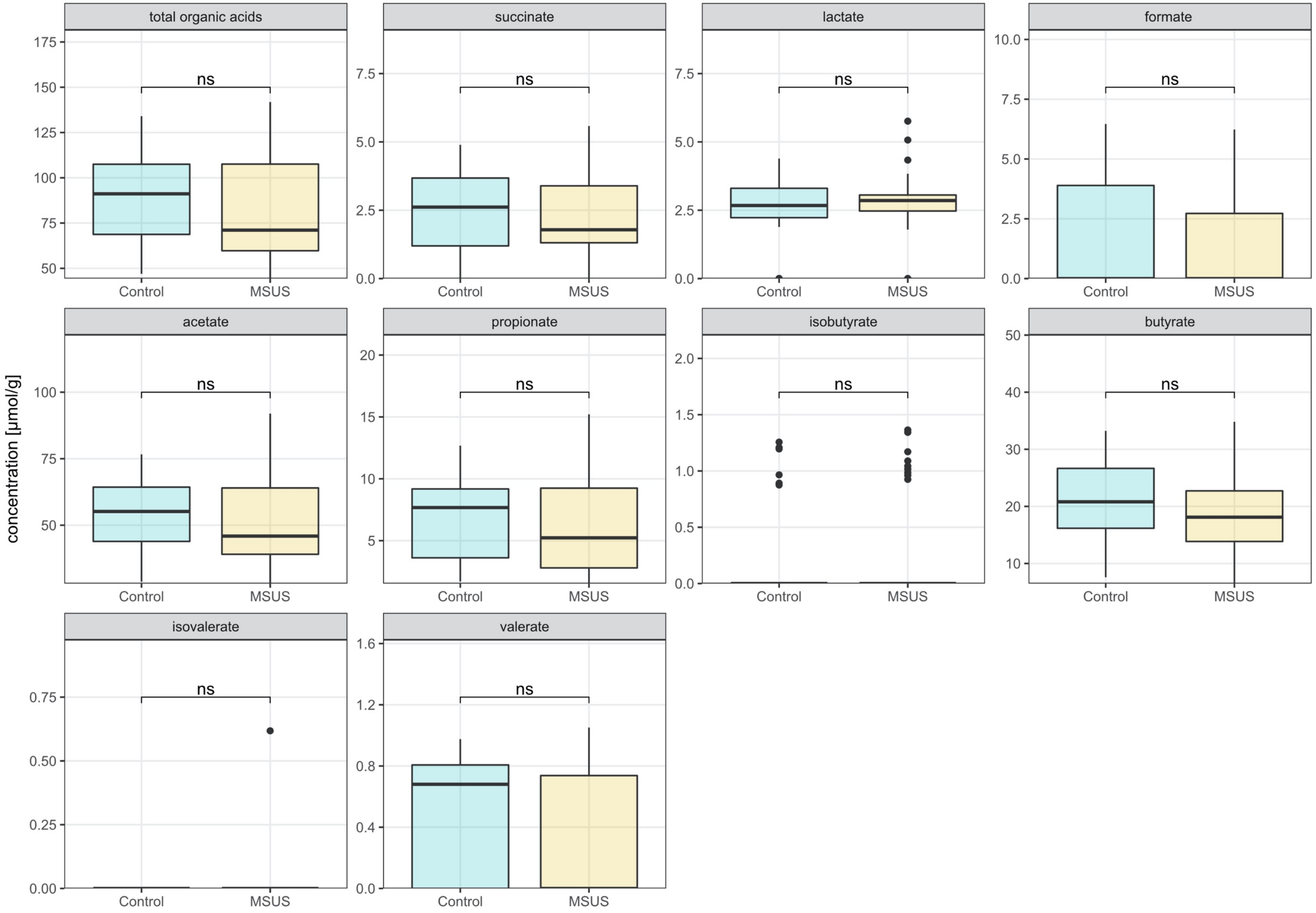


**Supplementary Figure 9: Effect of MSUS paradigm on cecal total and specific organic acid concentrations of 30-week-old F1 mice.** Comparison of MSUS *versus* control mice is displayed. Metabolite concentration is depicted per gram cecal wet weight. Boxplot with box elements showing upper and lower quantile and median. Whiskers extend from the hinge to ± 1.5 times the interquartile range or the highest/lowest value. Outliers are indicated as black points. Significances were calculated using generalized mixed effect models with FDR correction. ns: not significant

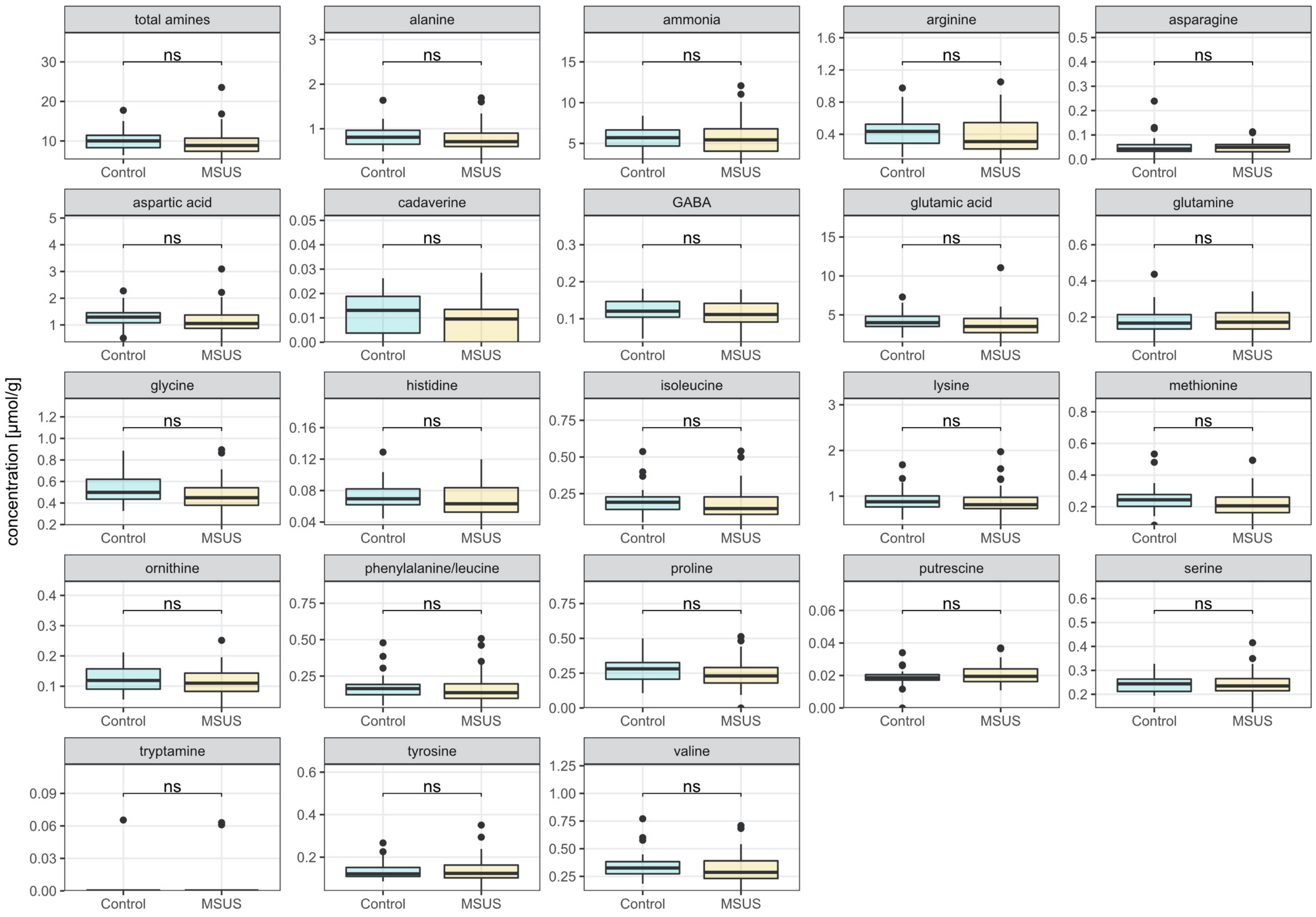


**Supplementary Figure 10: Effect of MSUS paradigm on cecal amine, amino acid, and ammonia concentrations of 30-week-old F1 mice.** Comparison of MSUS *versus* control mice is displayed. Metabolite concentration is depicted per gram cecal wet weight. Boxplot with box elements showing upper and lower quantile and median. Whiskers extend from the hinge to ± 1.5 times the interquartile range or the highest/lowest value. Outliers are indicated as black points. Significances were calculated using generalized mixed effect models with FDR correction. ns: not significant

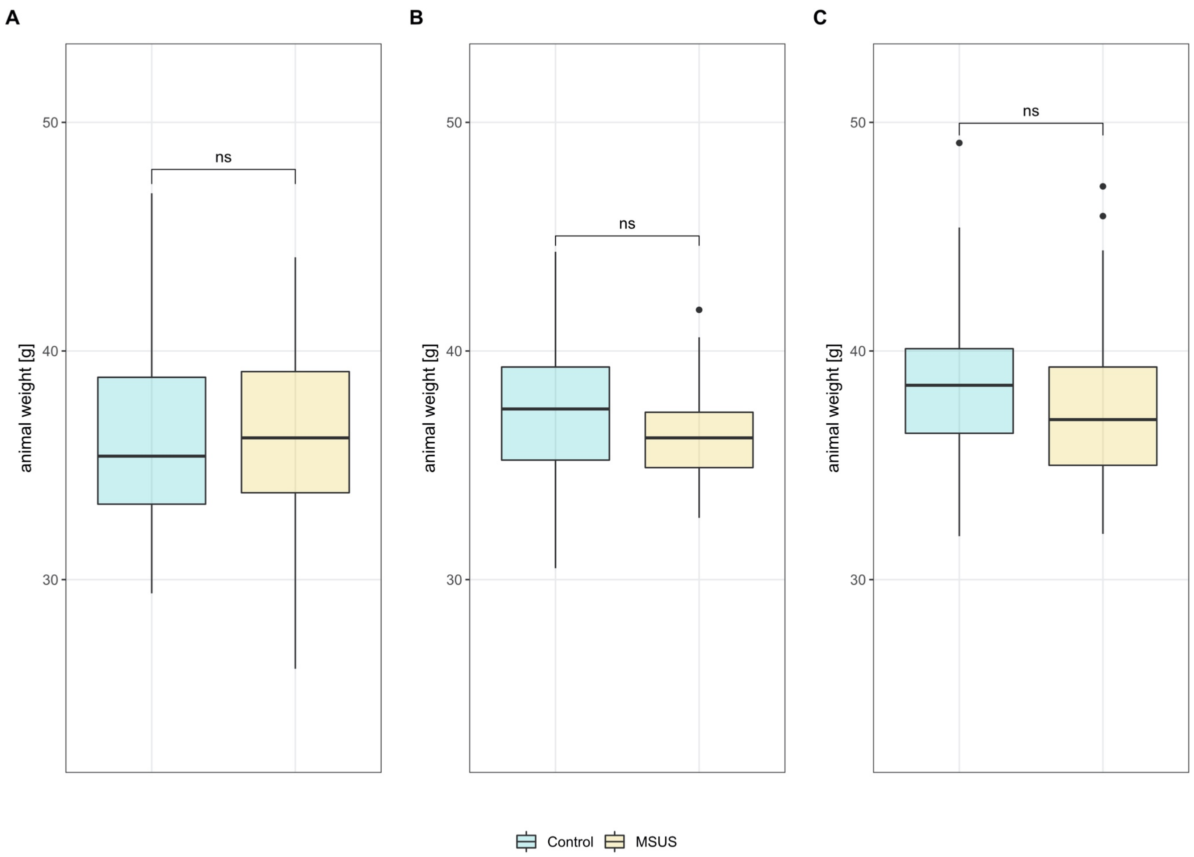


**Supplementary Figure 11: Effect of MSUS paradigm on animal weight of F1, F2, and F3 mice.** Comparison of MSUS *versus* control mice is displayed. Weight in gram of 30-week-old mice is depicted for (A) F1 and (C) F3, weight of 28-week-old mice is depicted for (B) F2. Boxplot with box elements showing upper and lower quantile and median. Whiskers extend from the hinge to ± 1.5 times the interquartile range or the highest/lowest value. Outliers are indicated as black points. Significances were calculated using generalized mixed effect models with FDR correction. ns: not significant

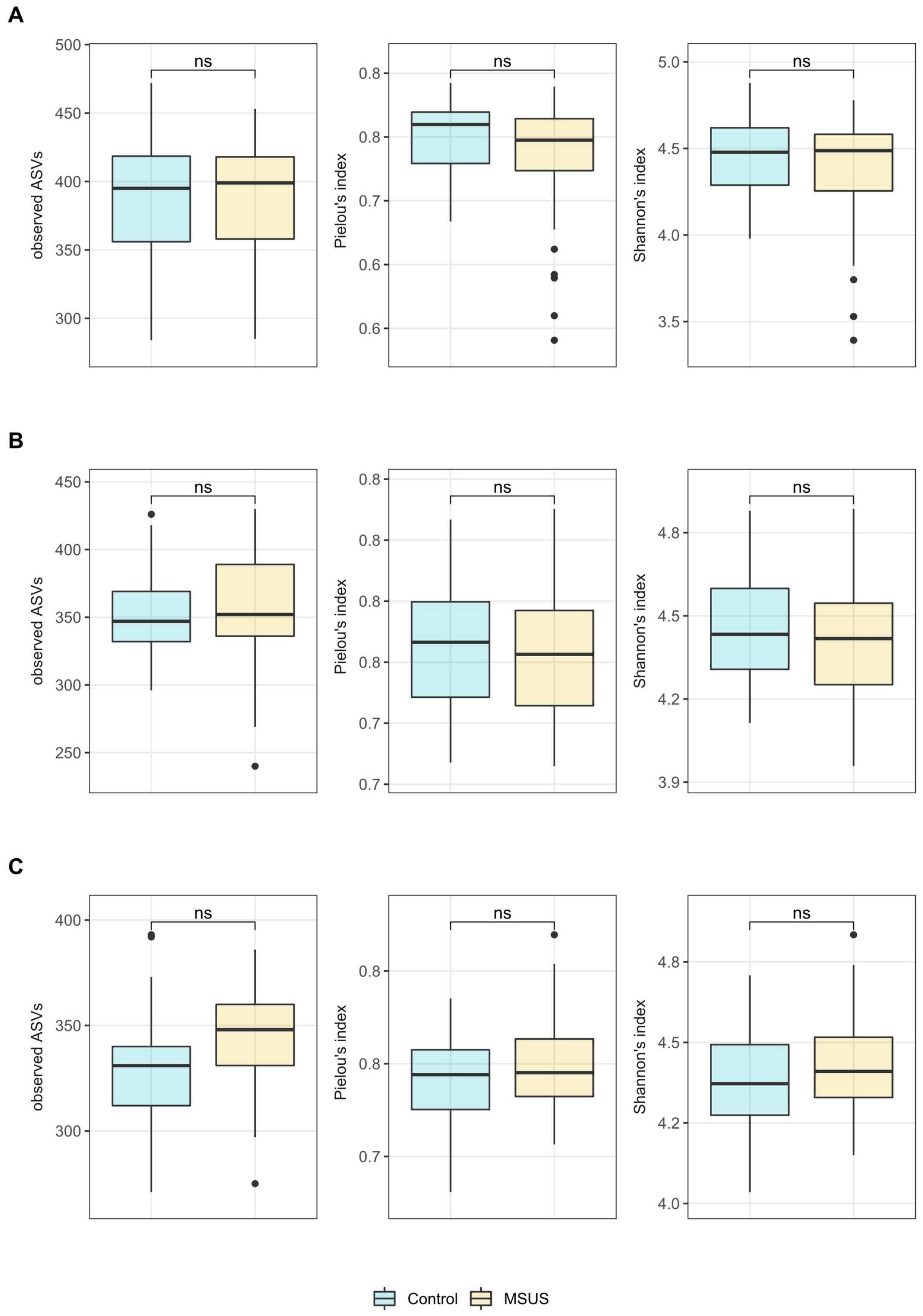


**Supplementary Figure 12: Effect of MSUS paradigm on alpha diversity metrics of F1, F2, and F3 mice fecal microbiota.** Comparison of richness (observed ASV), evenness (Pielou's index) and Shannon-diversity between MSUS and control is displayed. Alpha diversity of 30-week-old mice is depicted for (A) F1, and (C) F3, and alpha diversity of 28-week-old mice is depicted for (B) F2. Boxplot with box elements showing upper and lower quantile and median. Whiskers extend from the hinge to ± 1.5 times the interquartile range or the highest/lowest value. Outliers are indicated as black points. Significances were calculated using $\log_2$ transformed metrics and generalized mixed effect models with FDR correction. ns: not significant

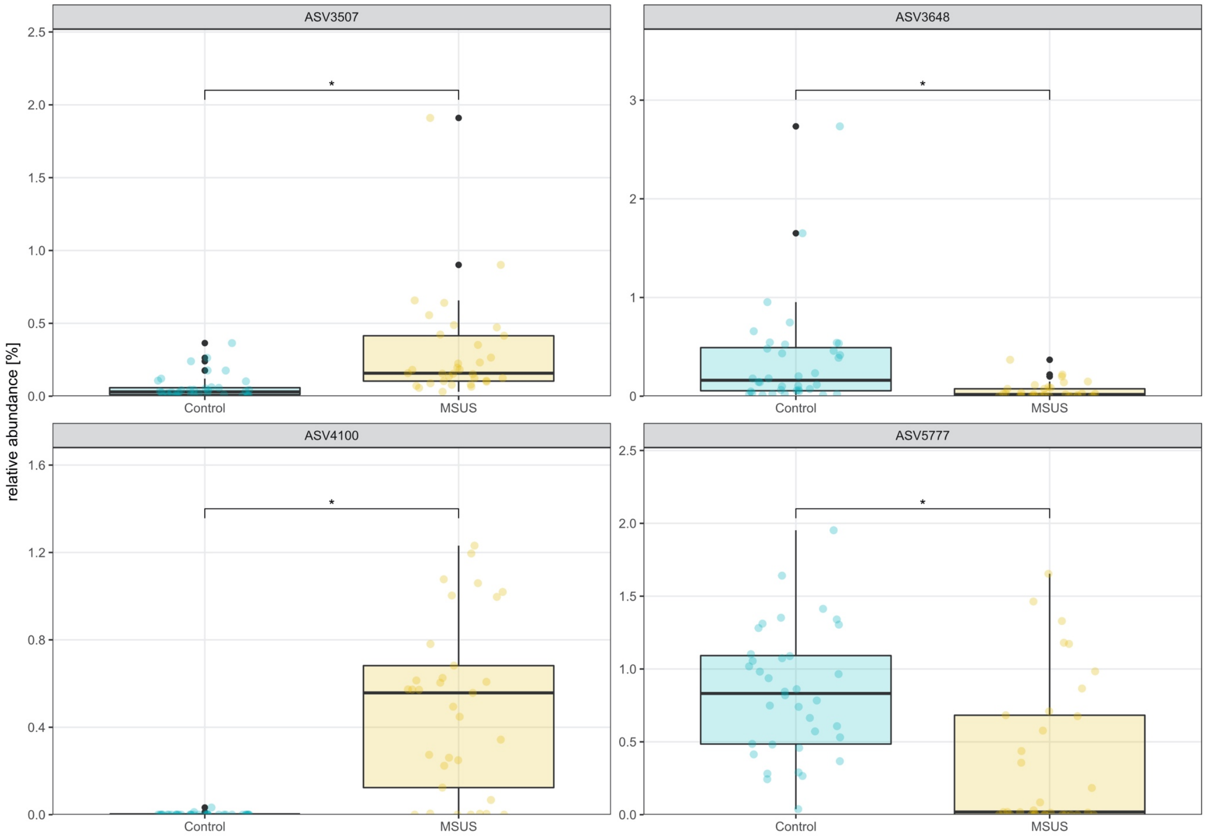


**Supplementary Figure 13: Effect of MSUS paradigm on relative abundance of ASVs in 28-week-old F2 mice fecal microbiota.** Significantly differentially abundant ASVs between MSUS *versus* control mice are displayed. Boxplot with box elements showing upper and lower quantile and median. Whiskers extend from the hinge to ± 1.5 times the interquartile range or the highest/lowest value. Outliers are indicated as black points, while colored points indicate individual microbiota. Significances were calculated using generalized mixed effect models with FDR correction. * $p < 0.05$

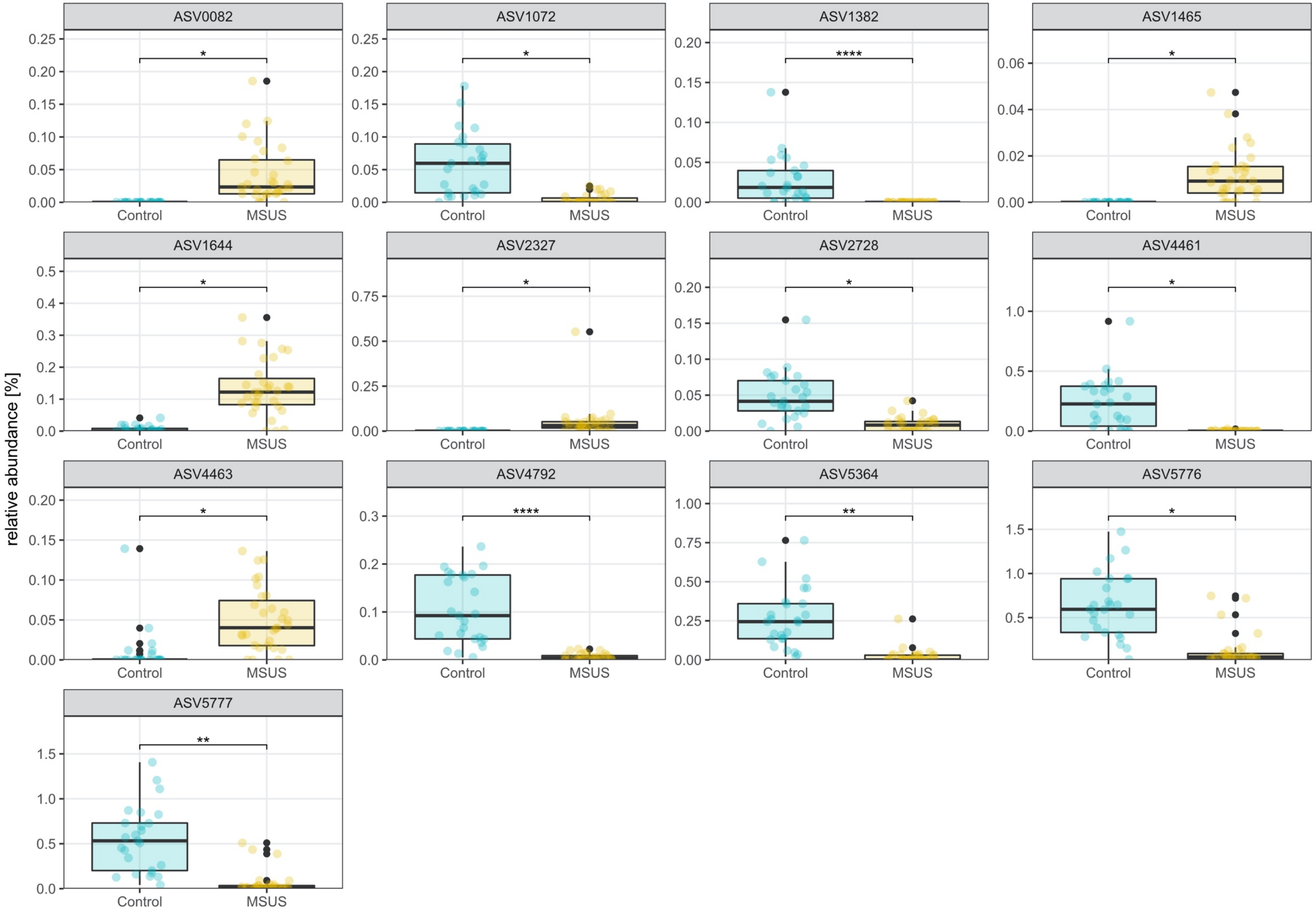


**Supplementary Figure 14: Effect of MSUS paradigm on relative abundance of ASVs in 30-week-old F3 mice fecal microbiota.** Significantly differentially abundant ASVs between MSUS *versus* control mice are displayed. Boxplot with box elements showing upper and lower quantile and median. Whiskers extend from the hinge to ± 1.5 times the interquartile range or the highest/lowest value. Outliers are indicated as black points, while colored points indicate individual microbiota. Significances were calculated using generalized mixed effect models with FDR correction. * $p < 0.05$; ** $p < 0.01$; **** $p < 0.0001$

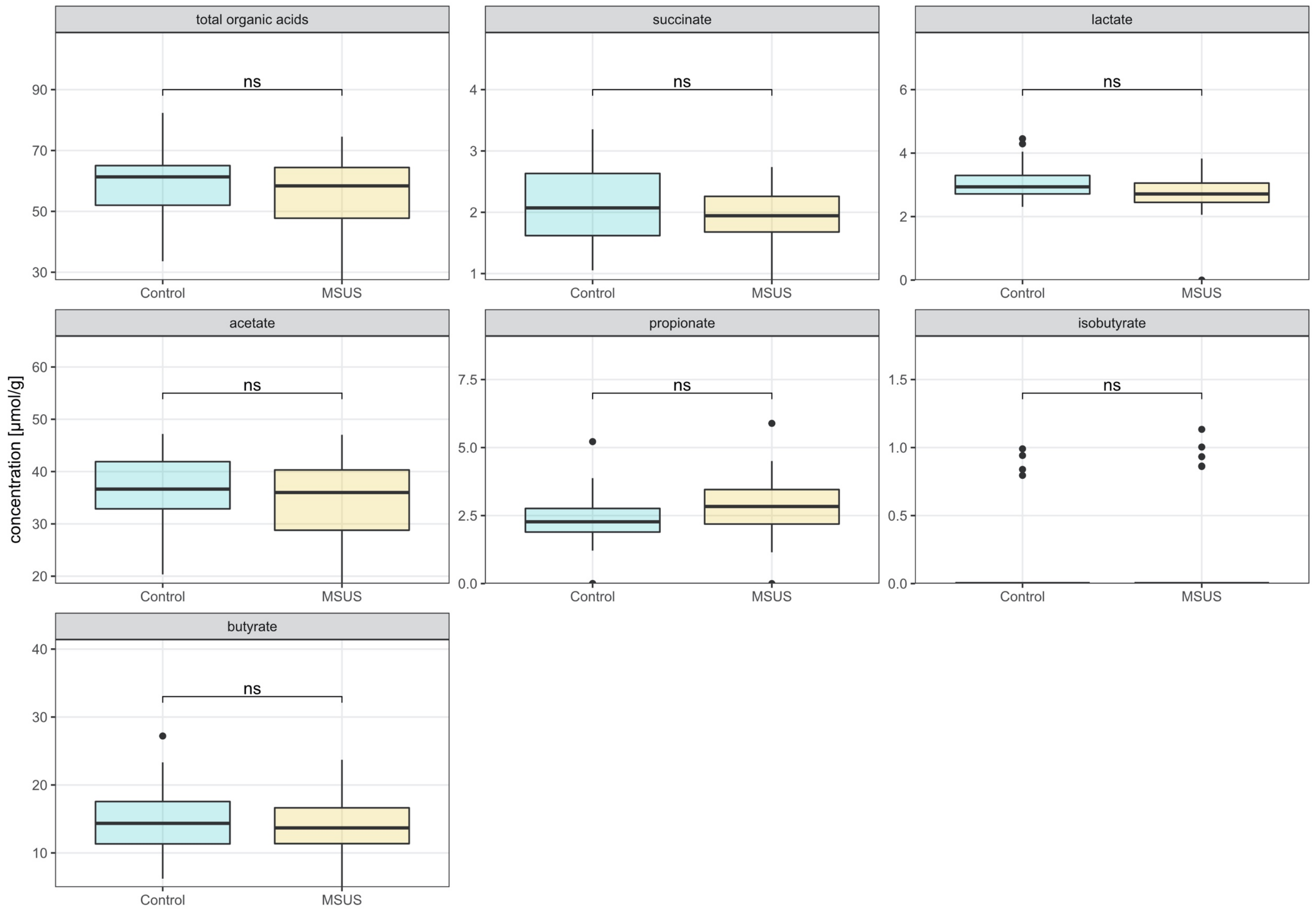


**Supplementary Figure 15: Effect of MSUS paradigm on cecal total and specific organic acid concentrations of 28-week-old F2 mice.** Comparison of MSUS *versus* control mice is displayed. Metabolite concentration is depicted per gram cecal wet weight. Boxplot with box elements showing upper and lower quantile and median. Whiskers extend from the hinge to ± 1.5 times the interquartile range or the highest/lowest value. Outliers are indicated as black points. Significances were calculated using generalized mixed effect models with FDR correction. ns: not significant

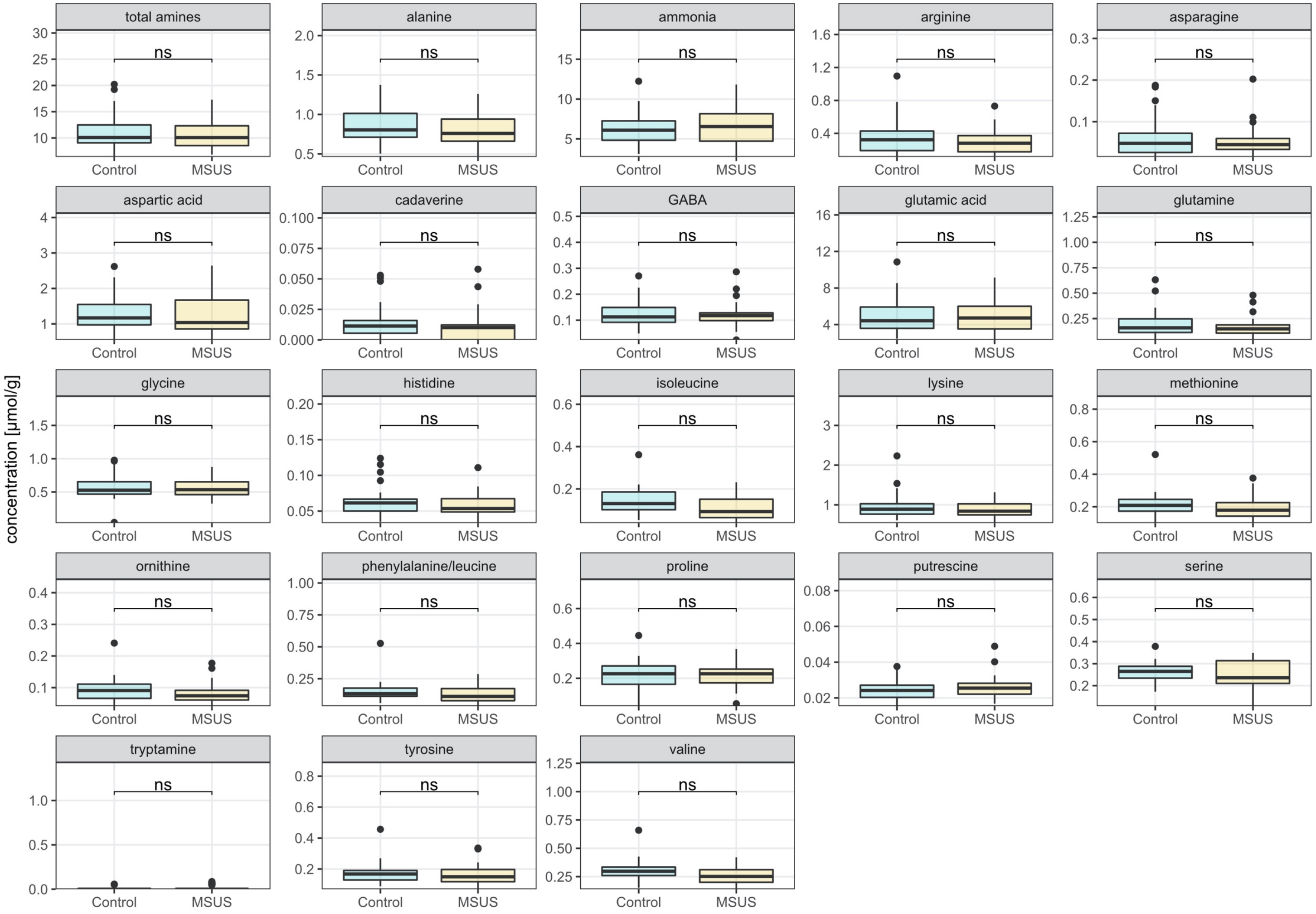


**Supplementary Figure 16: Effect of MSUS paradigm on cecal amine, amino acid, and ammonia concentrations of 28-week-old F2 mice.** Comparison of MSUS *versus* control mice is displayed. Metabolite concentration is depicted per gram cecal wet weight. Boxplot with box elements showing upper and lower quantile and median. Whiskers extend from the hinge to ± 1.5 times the interquartile range or the highest/lowest value. Outliers are indicated as black points. Significances were calculated using generalized mixed effect models with FDR correction. ns: not significant

**Supplementary Table 1: Statistical analysis of pairwise comparison of different beta diversity metrics across life span and between different phenotyping groups of control fecal microbiota.** Significances were calculated using PERMANOVA, and permutation test for dispersion, both including FDR correction**.**

| metric | comparison | *p*-value* | $R^2$ | dispersion *p*-value* |
|---|---|---|---|---|
| binary Jaccard index | PND30vsPND22 | 0.008 | 0.162 | 0.665 |
| | WEEK08vsPND22 | 0.006 | 0.205 | 0.825 |
| | WEEK15vsPND22 | 0.006 | 0.232 | 0.471 |
| | WEEK30vsPND22 | 0.004 | 0.216 | 0.665 |
| | WEEK08vsPND30 | 0.003 | 0.106 | 0.086 |
| | WEEK15vsPND30 | 0.003 | 0.120 | 0.011 |
| | WEEK30vsPND30 | 0.003 | 0.118 | 0.055 |
| | WEEK15vsWEEK08 | 0.045 | 0.056 | 0.196 |
| | WEEK30vsWEEK08 | 0.024 | 0.057 | 0.530 |
| | WEEK30vsWEEK15 | 0.016 | 0.059 | 0.825 |
| | GR2vsGR1 | 0.927 | 0.104 | 0.825 |
| weighted Jaccard index | PND30vsPND22 | 0.008 | 0.180 | 0.007 |
| | WEEK08vsPND22 | 0.006 | 0.196 | 0.031 |
| | WEEK15vsPND22 | 0.006 | 0.214 | 0.086 |
| | WEEK30vsPND22 | 0.004 | 0.195 | 0.018 |
| | WEEK08vsPND30 | 0.003 | 0.120 | 0.007 |
| | WEEK15vsPND30 | 0.003 | 0.122 | 0.022 |
| | WEEK30vsPND30 | 0.003 | 0.129 | 0.007 |
| | WEEK15vsWEEK08 | 0.006 | 0.064 | 0.972 |
| | WEEK30vsWEEK08 | 0.003 | 0.067 | 0.665 |
| | WEEK30vsWEEK15 | 0.003 | 0.076 | 0.665 |
| | GR2vsGR1 | 0.592 | 0.109 | 0.972 |

* FDR-adjusted; GR1: group 1 (behavioral phenotyping before breeding); GR2: group 2 (breeding before behavioral phenotyping); PND: postnatal day

**Supplementary Table 2: Statistical analysis of comparison of different beta diversity metrics between F1 MSUS and control fecal microbiota across life span.** Significances were calculated using PERMANOVA, and permutation test for dispersion, both including FDR correction**.**

| metric | age compared | *p*-value* | $R^2$ | dispersion *p*-value* |
|---|---|---|---|---|
| binary Jaccard index | PND22 | 0.8 | 0.188 | 0.964 |
| | PND30 | 0.002 | 0.063 | 0.710 |
| | WEEK08 | 0.002 | 0.061 | 0.964 |
| | WEEK15 | 0.002 | 0.067 | 0.964 |
| | WEEK30 | 0.002 | 0.060 | 0.710 |
| weighted Jaccard index | PND22 | 0.119 | 0.209 | 0.014 |
| | PND30 | 0.002 | 0.058 | 0.870 |
| | WEEK08 | 0.119 | 0.052 | 0.979 |
| | WEEK15 | 0.075 | 0.055 | 0.710 |
| | WEEK30 | 0.119 | 0.051 | 0.964 |

* FDR-adjusted; PND: post-natal day

**Supplementary Table 3: Statistical analysis of comparison of different beta diversity metrics between MSUS and controls of F1, F2, and F3 fecal microbiota.** Significances were calculated using PERMANOVA, and permutation test for dispersion, both including FDR correction**.**

| **metric** | **age compared** | ***p*-value*** | **$R^2$** | **dispersion *p*-value*** |
|---|---|---|---|---|
| binary Jaccard index | F1 (WEEK30) | 0.004 | 0.060 | 0.632 |
| | F2 (WEEK28) | 0.004 | 0.077 | 0.632 |
| | F3 (WEEK30) | 0.004 | 0.131 | 0.868 |
| weighted Jaccard index | F1 (WEEK30) | 0.101 | 0.051 | 0.868 |
| | F2 (WEEK28) | 0.048 | 0.071 | 0.632 |
| | F3 (WEEK30) | 0.011 | 0.099 | 0.868 |

* FDR-adjusted

**Supplementary Table 4: Time point and number of fecal samples collected for F1, F2, and F3 control and MSUS mice.**

| generation | treatment | age | mice | cages | litters |
|---|---|---|---|---|---|
| F1 | control | PND22 | 11 | 3 | 3 |
| | MSUS | PND22 | 12 | 3 | 3 |
| | control | PND30<br>8-weeks-old<br>15-weeks-old | 37 | 11 | 13 |
| | MSUS | PND30<br>8-weeks-old<br>15-weeks-old | 50 | 12 | 11 |
| | control | 30-weeks-old | 35 | 11 | 13 |
| | MSUS | 30-weeks-old | 49 | 12 | 11 |
| F2 | control | PND21 | 20 | 4 | 4 |
| | MSUS | PND21 | 25 | 6 | 6 |
| | control | 16-week-old | 36 | 9 | 13 |
| | MSUS | 16-week-old | 36 | 10 | 14 |
| | control | 28-weeks-old | 36 | 9 | 14 |
| | MSUS | 28-weeks-old | 33 | 9 | 14 |
| F3 | control | 30-weeks-old | 25 | 7 | 8 |
| | MSUS | 30-weeks-old | 31 | 8 | 8 |

F1: first generation; F2: second generation; F3: third generation; PND: postnatal day
